# Exogenous Methylglyoxal Improves Maize Yield by Alleviating Plant Diabetes and leaf senescence under Drought

**DOI:** 10.1101/2023.07.18.549499

**Authors:** Yi-Hsuan Lin, Ya-Ning Zhou, Yu-ka Jin, Zu-Dong Xiao, Ying-Jun Zhang, Cheng Huang, Bo Hong, Zhen-Yuan Chen, Si Shen, Shun-Li Zhou

## Abstract

Drought-induced leaf senescence is related to high sugar levels in leaves, photosynthesis inhibition, and ultimate yield loss. This physiological phenomenon in leaves bears resemblance to the symptom of diabetes in human disease. However, the underlying mechanisms of plant diabetes on carbon imbalance in maize leaf and corresponding detoxification strategy have not been well understood. In this study, we demonstrated that foliar application of exogenous methylglyoxal (MG) delayed leaf senescence and promoted photoassimilation, retrieved 14% yield loss induced by drought stress during grain filling stage. Transcriptome and metabolite analysis revealed that drought increased sugar accumulation in leaf with inhibition of sugar transporters facilitating phloem loading. This further lead to disequilibrium of glycolysis and over-accumulation of endogenous MG. Contrarily, exogenous MG significantly upregulated glycolytic flux and glyoxalase system catabolizing endogenous MG and advanced glycation end products toxicity, ultimately alleviating plant diabetes. Besides, the genes facilitating anabolism and catabolism of trehalose- 6-phosphate were promoted and suppressed by drought, respectively, whereas exogenous MG reversed the effect. Moreover, exogenous MG activated phenylpropanoid biosynthetic pathway, likely promoting cell structural integrity. Collectively, these results suggest that exogenous MG alleviates the toxic effect from drought-induced sugar accumulation and activates the defense-related pathway, thereby maintaining leaf function and yield production.

**Highlight:** Exogenous methylglyoxal stimulates glycolytic flux and glyoxalase system, providing a potential insight to alleviate plant diabetes under drought condition.

## Introduction

With the increase of extreme climate events, food security was challenged by abiotic stresses (FAO, 2021). Among them, drought stands out as the most devastating event in crop production, caused an annual economic loss around 3.7 billion worldwide (FAO, 2021). Moreover, the intensification and frequent occurrence of flash drought events worldwide have posed significant challenges to drought forecasting and mitigation capabilities (Yuan et al., 2023). Maize, one of the most important cereals, is with versatile utilization, including food supply for human diets, forage for livestock, and raw materials for biofuel production (Dietz et al., 2021). However, according to environmental policy impact climate model, the risk of global drought stress leading to maize yield reduction at the end of 21^st^ century will increase by 19.18% (Guo et al., 2016). In this case, drought has detrimental effect on stay-green and function of leaves, and thus fundamentally limits yield and yield potential, particularly during flowering and grain filling stages of maize production (NeSmith and Ritchie, 1992; Shen et al., 2020).

Leaf senescence is accelerated in response to drought, which further inhibits the supply of photoassimilate to kernels for yield formation (Jurgens et al., 1978). Notably, sugar is a crucial factor in inducing leaf senescence (Van Doorn, 2008). The initiation step of drought-induced leaf senescence is overaccumulation of sugars in leaf cells, which is prior to chlorophyll degradation (Van Doorn, 2008). Moreover, studies have shown that exogenous sugars can induce senescence in intact leaf through sugar signaling and phytohormones signaling (Asim et al., 2022b; Asim et al., 2022c). Accordingly, the theory of ‘plant diabetes’ was proposed to describe the symptom of sugar overaccumulation in cells that inevitably leads to toxic sugar derivatives and thus damages plant cells (Shimakawa et al., 2014; Takagi et al., 2014; Rivero-Marcos and Ariz, 2022). Plant diabetes is a conserved and universal phenomenon that causes dysfunction of leaves under various stresses such as NH4^+^, osmotic, heat, and high light stresses (Paudel et al., 2016; Borysiuk et al., 2018; Chaplin et al., 2019). Nonetheless, how to detoxify the plant diabetes in leaf senescence and further alleviate yield penalty in coping with stresses such as drought has received limited attention.

Methylglyoxal (MG) is the main sugar derivative that is generated from glycolysis and indirectly regulated by triose phosphate isomerase (TPI). MG accumulation can cause toxic damage to cell functioning and proliferation (Chen and Thelen, 2010). MG alters the structures and stabilities of amino acids and nucleic acids, leading to the formation of advanced end glycation products (AGEs), in terms of protein carbonylation (Allaman et al., 2015; Li et al., 2017; Fu et al., 2021). Similarly, MG- derived AGEs are also associated with the complication of human diseases such as diabetes, obesity, and aging (Mostofa et al., 2018). To maintain MG homeostasis, glyoxalase system consisting of glyoxalase I (Gly I) and glyoxalase II (Gly II) facilitates detoxification of MG in organisms (RACKER, 1951; CROOK and LAW, 1952). Even though, endogenous MG usually accumulates and escalates to cause dysfunction of plant cells or tissues under various stresses (Singla-Pareek et al., 2020). Intriguingly, instead of being toxic, MG plays a dual role in regulation of signal transduction with a dose dependent manner. As a signal initiator, MG was associated with regulation of protein kinase and transcription factor in yeast and rice, respectively (Maeta et al., 2005; Kaur et al., 2015). Recently, it has been reported that exogenous MG possibly affected the homeostasis of endogenous MG under drought condition (Lin et al., 2022). However, it remains unclear how exogenous MG affects plant diabetes and underlying mechanisms of drought tolerance in maize.

In this study, we aim to (i) investigate how plant diabetes is involved in leaf senescence under drought condition. (ii) To understand the impact of application of exogenous MG on leaf senescence, yield formation, and physiological basis of MG- mediated tolerance to drought. By conducting RNA-sequencing, enzymatic activity, and metabolites analysis, we overview these key responses in order to decipher MG’s role in regulating source strength and yield formation of maize under drought stress. These results provide theoretical basis and practical approach to maintain or improve crop yield in face of global environmental challenge.

## Materials and methods

### Plant materials and arrangement

The experiment was conducted at Wu Qiao Experimental Station of China Agricultural University in Hebei province, China. Maize variety Zhengdan 958, a prevalent and generally used hybrid, was selected in this experiment. The density of maize cultivation was 56,000 per hectare with basal compound (500 kg ha^−1^; N 15%, P_2_O_5_ 15%, K_2_O 15%) at seedling stage and a top dressing of urea (165 kg ha^−1^; N 46%) at the V13 stage. Pesticide was used appropriately that there were no severer insect pests or diseases during whole growing season.

### Drought simulation, exogenous MG application, and sampling

To simulate the drought condition in the field circumstance, maize plants were grown in 20 outdoor pools with 10.98-m^2^ areas and 0.6-m depth for each pool (Fig. 1A). Drought pools have a movable canvas roof to prevent rainfall. For precise monitoring of the extent of drought stress (DS), artificially quantifying irrigation was conducted based on continuous monitoring of soil water content throughout the experiment using soil moisture sensor (Thetaprobe ML2x, Delta-T Device, UK). Soil volume water content of 20-30 cm depth was detected at 3, 6, 9, 12, 15, 18, 21, 24, 27, 29, and 30 DAP (Fig. 1B). According to our previous studies (Shen et al., 2020b; Chen et al., 2022), until the soil volume water content decreased to ∼35%, the maize plant demonstrated the drought effects of kernels developmental stunted and leaf senescence. Therefore, we defined 15 DAP as the drought commencement until 30 DAP, which was correspondence to 1-15 days after drought induction (DAI). At 30 DAP, pools were re-watered and maintained the well-watered status (∼60%) until maturity. In the control, pools were well-watered and maintained the soil volume water content at 60% on average until maturity (Fig. 1B).

**Fig. 1.**
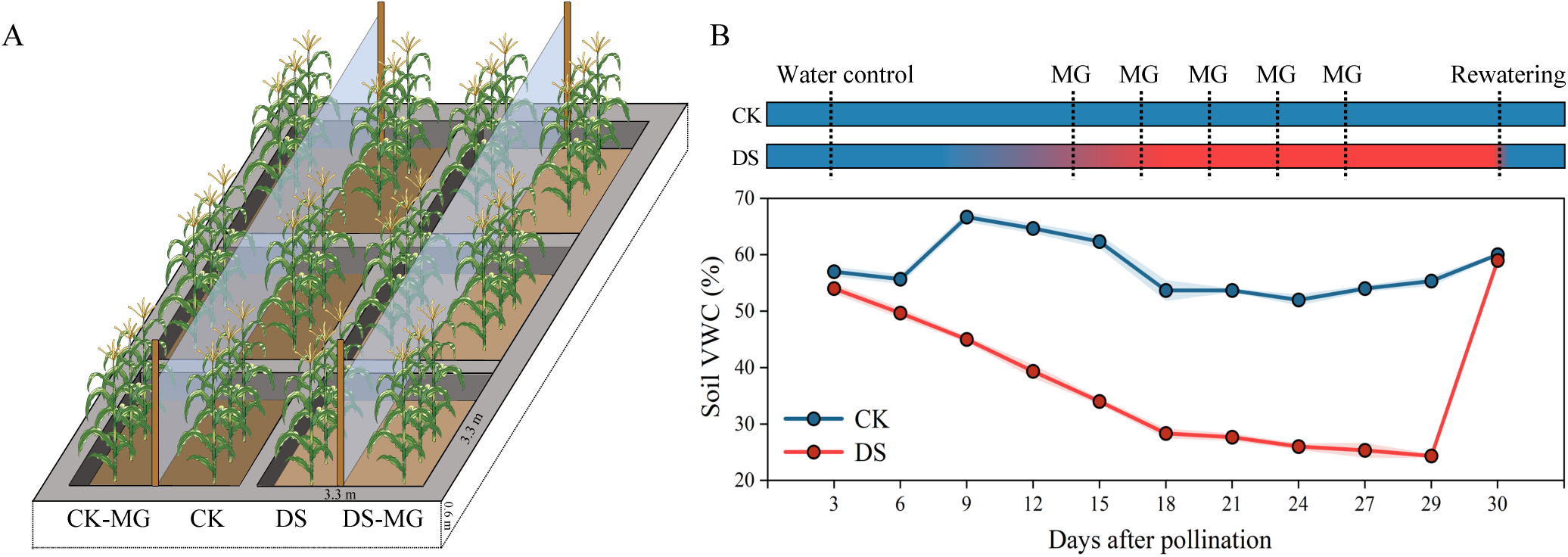
(A) Schematic diagram of drought pools, each pool has divided by transparent polyethylene sheet. (B) Water dynamic during well-watered and drought with the timeline of exogenous MG treatment. The shading area of blue and red indicated the error bar with the means (± SE) under CK and DS, respectively. Soil volume water content was measured in 20-30 cm depth at 3 days intervals. CK, well-watered; DS, drought stress.

Each pool would divide into two sections by placing the transparent polyethylene sheet (3 m X 2 m) (Fig. 1A). At 14, 17, 20, 23, and 26 DAP, 25 mM MG solution (Macklin) was applied in each treatment by foliar spraying and distilled water was used as the control (Fig. 1B). The details of spraying method have been described in previous study (Lin et al., 2022).

At 18, 21, 24, 27, and 30 DAP, correspondence to 3, 6, 9, 12, and 15 DAI, middle part of ear leaf was collected at 10 am and immediately frozen by liquid nitrogen stored in ultra-low freezer for the further physiology and biochemistry determination. Yield and yield components were measured at maturity, kernel sample was determined after 48 hr of drying at 80°C in oven.

### Measurement of SPAD value, chlorophyll fluorescence

With a view to evaluating the state of leaf senescence, relative chlorophyll content was measured from the lower part of leaf to the top of leaf (7^th^ to 20^th^). Each leaf has measured for 20 spots to represent the relative chlorophyll content by SPAD meter (Chlorophyll meter SPAD-502Plus, Japan). Chlorophyll fluorescence (Fv/Fm and YII) was measured at ear leaf after 30 minutes dark adaption by Walz (MINI-PAM-II, Germany). SPAD value and chlorophyll fluorescence determination were carried out at 3, 6, 9, 12, and 15 DAI.

### Measurement of glucose, fructose, sucrose, and starch content

Middle part of ear leaf was used for extraction of sugars. According to (Hendrix, 1993), fresh samples 0.1 g were ground into powder by liquid nitrogen, added 1.5 mL of 80% ethanol and heat up in 80°C water for 30 min. After heating up, centrifuged at 12000 g for 5 min and repeated the step above twice until the soluble sugar was completely extracted. The remains left in the tube were used for starch extraction. Added 0.5 mL distilled water, boiled in water for 10 min. Added 1 mL of 200 mM sodium acetate buffer (10 mM CaCl_2_, 250 U amyloglucosidase, 20U α- amylase, at pH 6), reaction in 60°C for 6 hr. The samples were centrifuged at 10000 g for 5 min. The supernatant was prepared for the following determination (Smith and Zeeman, 2006; Sørensen et al., 2018). Before the determination of sugar content, each sample has added enough charcoal to decolorize the pigments.

Glucose, fructose, sucrose, and starch content were determined by investigating the generation of NADH at 340 nm. First, glucose and starch content were added 10 µl of samples and 190 µl of mixture I which contained 100 mM HEPES at pH 7.5, 4 mM MgCl, 1 mM NAD^+^, 1.2 mM ATP. Afterward, 5 µl of mixture II was added, which contained 1.5 U hexokinase, 1.2 U glucose-6-phosphate dehydrogenase, and reaction for 40 min at room temperature. Fructose content was conducted after glucose content determination. Added 5 µl of 0.04 U/µl glucose-6-phosphate isomerase to the former wells and completely reacted for 40 min. Sucrose content was determined by adding 5 µl of 9 U invertase to former wells and completely reacted for 100 min. Quantification of sugar content was calculated with standard curve.

### Determination of glycolytic enzymes activity

Measurement of activity for phosphofructokinase and glyceraldehyde-3-phosphate dehydrogenase was extracted and estimated by FPK essay kit and GAPDH easy kit (Grace biotechnology, China), respectively. Triosephosphate isomerase easy has been described by (Sharma et al., 2012) with little modification. Homogenate of fresh sample 0.1 g was added 1 mL extracted buffer containing 50 mM Tris-HCl (pH 8.0), 1 mM EDTA, 10% (v/v) glycerol, 5 mM dithiothreitol, and 5 % PVP. The homogenates were centrifuged for 15 min at 12000 g at 4°C and diluted for 5 times in extracted buffer. Added the mixture contained 0.1 M Tris-HCl (pH 7.5), 1 mM glyceraldehyde-3- phosphate, 0.5 mM EDTA, 2.5 U glycerophosphate dehydrogenase, and 0.2 mM NADH to start the reaction. The activity estimation was performed by NADH disappearance overtime at 340 nm.

Determination of MG, AGEs adduct content, and glyoxalase system activity Extraction of MG content was followed by (Mustafiz et al., 2010), fresh sample 0.1 g was homogenized in 1 mL of 0.5 M perchloric acid and incubated on ice for 10 min. Homogenates were centrifuged at 15,000 g for 20 min at 4 °C. 10 mg of charcoal was added in the supernatant for decolorized and neutralized sample with 1 M CK_2_O_3_. The sample was centrifuged again for the clear supernatant. MG quantification was conducted by adding 7.2 mmol 1, 2-diaminobenzene after 30 min and investigated at 336 nm. MG content was calculated with standard curve and expressed as µmol g^−1^ FW. Methylglyoxal hydroimidazolone (MG-H1) adduct and carboxymethyl-lysine (CML) adduct was extracted by using butyl alcohol: methanol: water (5: 25: 70 v: v: v) and performed in MG-H1 ELISA kit and CML ELISA kit (Shanghai xinyu biotechnology, China), respectively. Glyoxalase system essay was performed according to (Mustafiz et al., 2010). Briefly, Gly I and Gly II crude protein were extracted from fresh sample 0.25 g by buffer contained 50 mmol Tris–HCL (pH 7.0), 16 mmol MgSO_4,_ 0.2 mmol phenylmethanesulfonyl fluoride, and 0.2% PVP-40. The activity of Gly I and Gly II were estimated the formation and broke down of SLG at 240 nm, respectively.

### In vitro leaf strip senescence assay

Leaf strips of 2 cm in length were detached from middle part of 5^th^ leaf from ZD958 at V5 stage and were kept in 50 mM mannitol (as control), glucose, fructose, and sucrose with or without adding 1 mM N-acetyl-L-cysteine (NAC, MG scavenger, (Li et al., 2018)) for 4 days. The chlorophyll content was detected spectrophotometrically after extraction in 95% ethanol and calculated by the formula: Total chlorophyll content= 5.24*A664.2+ 22.24*A648.6 (Lichtenthaler, 1987). Four biological replicates and three technical replicates were performed in the experiment.

### Total RNA extraction, library construction, and transcriptome sequencing

Total RNA was extracted from middle part of ear leaf at 12 DAI with the RNA extraction Kit (DP411, TIANGEN, China). RNA purity, concentration, and integrity were detected by the NanoDrop2000 Spectrophotometer (Thermo Fisher, US) and Agient2100/LabChip GX (PerkinElmer, US), respectively. Qualified samples were used for constructing the transcriptome library by PCR product enrichment. The libraries were qualified by Qsep-400 then sequenced on an Illumina NovaSeq6000 platform (Illumina, US). The expression of each gene was transformed to transcript per Million (TPM) to compare different treatments.

### Differential gene expression and bioinformatics analysis

The filtered clean data was aligned to the reference genome of Zea_mays. AGPv4.40 by HISAT2 and estimated the differential gene expression between DS vs CK, CK-MG vs CK, and DS-MG vs DS, respectively. Genes with q ≤ 0.05 and fold change ≥ 1.5 were defined as differentially expressed genes (DEGs). Metabolic pathway enrichment analysis of DEGs were performed by using gene ontology (GO) and Kyoto Encyclopedia of Genes and Genomes (KEGG) in BMKCloud platform (http://www.biocloud.net). The p–value ≤ 0.05 were defined as significantly enriched. Gene IDs involved in metabolic pathways were aligned and identified to MaizeGDB (https://www.maizegdb.org).

### Statistical analysis

All measurements above included at least three to five biological replications from individual plants. Student’s t test and one-way ANOVA using Duncan’s new multiple range test were conducted in the data. Experimental data was analyzed by IBM SPSS Statistics, Microsoft Excel, and R Studio. Figures were illustrated by Microsoft PowerPoint, OriginPro, Adobe Fresco, and R Studio.

## Results

### Exogenous MG retrieved yield loss under drought

The ear phenotypes at 15 DAI (the end of drought conduction) and at maturity were estimated to represent the drought and both MG effects. At 15 DAI, the ear length was shrunk, and the development of apical kernel was inhibited, by DS (Fig. 2A). However, compared to control, the ear length and development of kernel were enhanced by exogenous MG under DS, but there was no clear difference under well- watered (Fig. 2A). We further estimated the drought and both MG effects on yield and yield components at maturity. The ear was evenly divided into 10 sections based on the fertilized ovaries number, and each section’s kernel weight was determined on average (Fig. 2A-C). The weight of apical kernel from sections 7-10 was significantly decreased by 0.2-13 times under DS, compared to well-watered (Fig. 2B). Exogenous MG significantly increased the weight of apical kernel (from sections 7-10) by 0.1-8 times under DS, compared to control (Fig.2B). Moreover, the abortion number and yield under DS were significantly higher by 5.5 times and lower by 0.19 times, respectively, than well-watered (Fig. 2D, E). Exogenous MG significantly decreased abortion number and retrieved yield loss by 64% and 14% under DS, respectively, compared to control (Fig. 2D, E). Our data reveal that exogenous MG enhanced the drought tolerance by reducing yield penalty under DS.

**Fig. 2.**
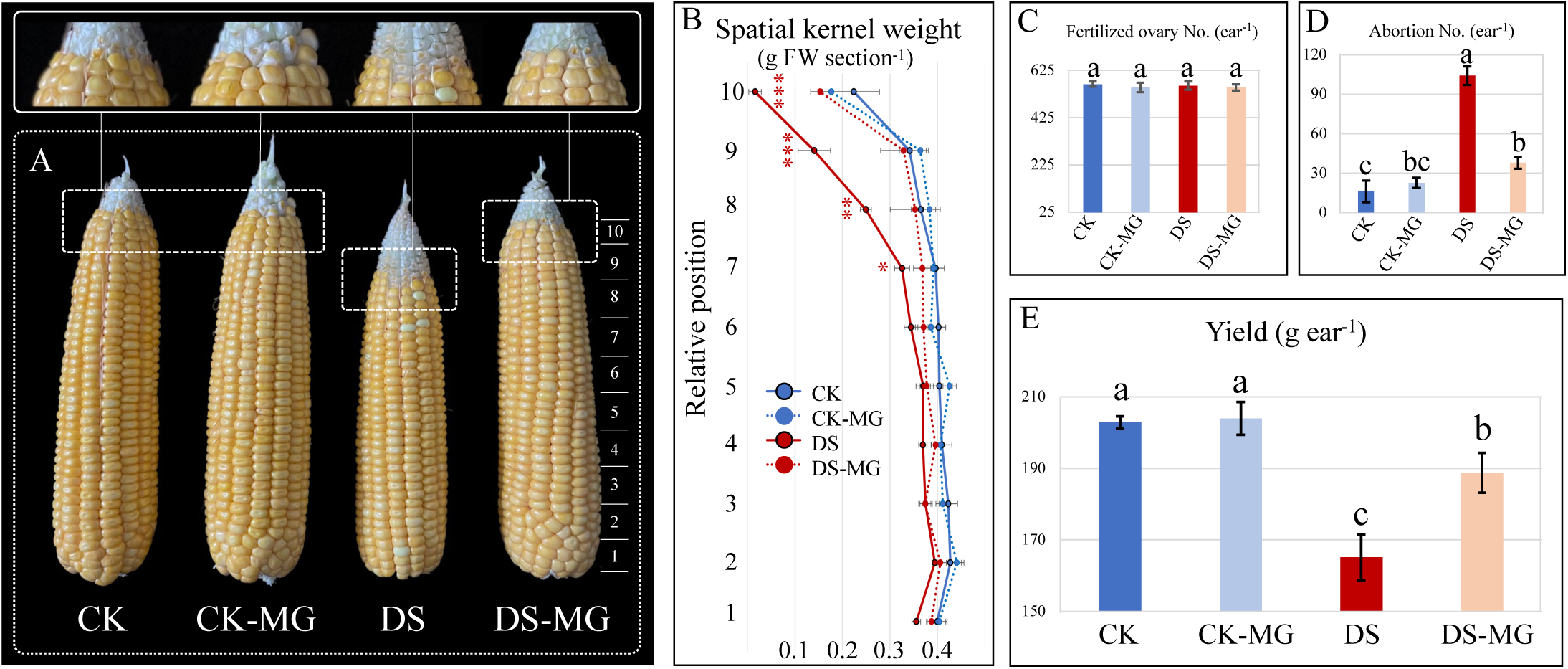
The effects of drought and exogenous MG on (A) phenotype of ear at 15 DAI, (B) spatial kernel weight, (C) fertilized ovaries number, (D) kernel abortion number, and (E) yield under well-watered and drought conditions at maturity. CK, well- watered; DS, drought stress; DAI, days after drought induction. Data are means (± SE). Asterisks indicate significant difference between DS-MG and DS (t test, n=5, *p<0.05, **p<0.01, ***p< 0.001). One-way ANOVA was conducted using Duncan’s new multiple range test, and different letters above the bars indicate significant differences (p < 0.05).

### Exogenous MG delayed drought-induced leaf senescence and improved photosynthetic capacity

To investigate the direct effects of exogenous MG on source capacity as drought progressed, the dynamics of leaf’s SPAD value and chlorophyll fluorescence were analyzed (Fig. 3). At 15 DAI, the lower part of leaf under DS exhibited severe curling and color-fading compared to well-watered (Fig. 3B). Visually, exogenous MG delayed the drought-induced senescence of leaves from lower position of plants (Fig. 3B). The SPAD values across the plant (7^th^-20^th^) under DS indicated that the sequence and extent of leaf senescence from lower part to top and the ear part were more prominent compared to well-watered (Fig. 3C). Furthermore, the SPAD value of ear leaf under DS was significantly decreased by 6-8% from 12-15 DAI compared to well- watered (Fig. S.1). Exogenous MG under DS maintained the higher SPAD value in leaves across the plant from 6-15 DAI compared to control (Fig. 3C). Moreover, under DS, the Fv/Fm was significantly decreased by 4-8% from 12-15 DAI, and the YII was significantly decreased by 17% at 15 DAI compared to well-watered (Fig. 3D, E). Exogenous MG under DS significantly maintained higher Fv/Fm by 4-6%, and higher YII by 20-40% at 12-15 DAI, compared to control (Fig. 3D, E). Similarly, compared to control, exogenous MG significantly increased the YII by 14-26% from 12-15 DAI under well-watered (Fig. 3D, E). Overall, the data indicates that exogenous MG delayed the senescence all leaves of plant and improves the photosynthetic capacity under DS.

**Fig. 3.**
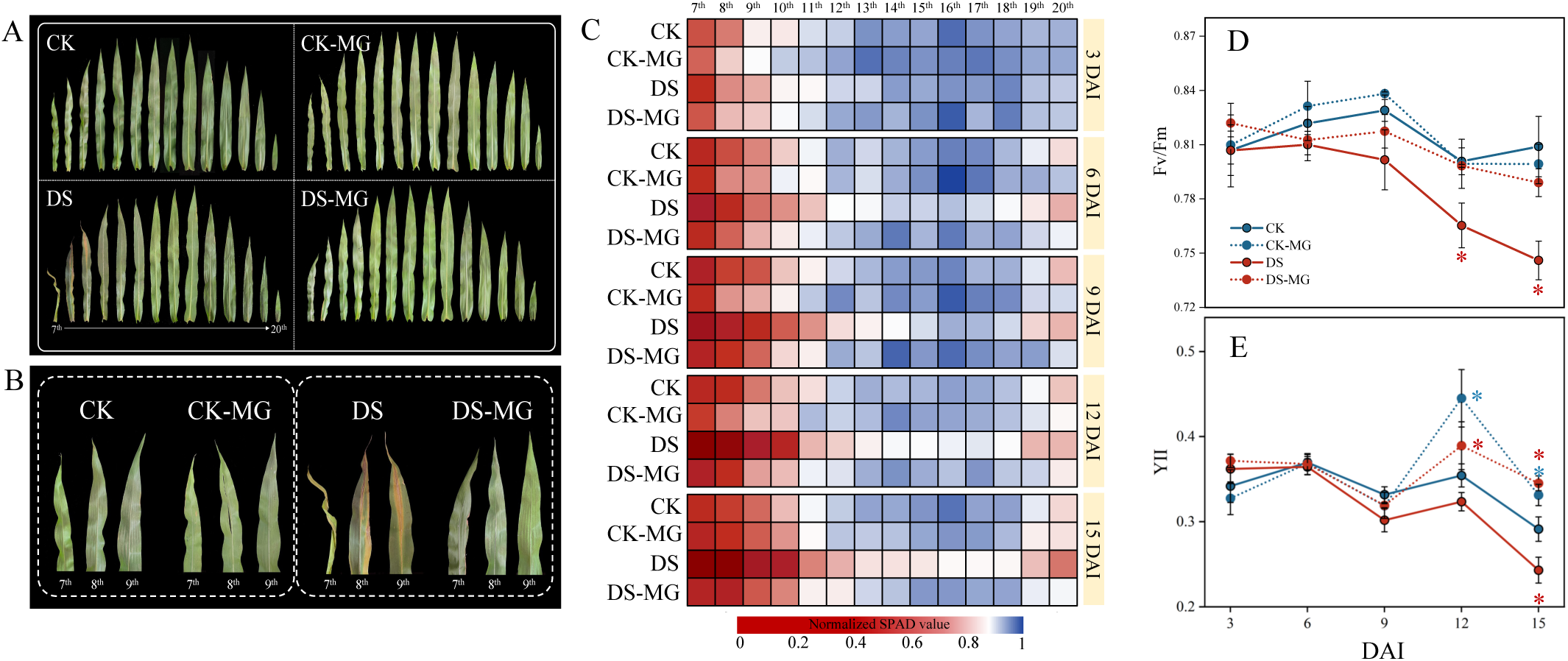
The effects of drought and exogenous MG on (A) phenotype of whole leaves and (B) lower part of leaves (7 -9) at 15 DAI, (C) normalized SPAD values of leaf from 7 to 20, and (D-E) photosynthetic capacity of ear leaf (15) under well-water and drought. CK, well-watered; DS, drought stress; DAI, days after drought induction. Data are means (± SE). Asterisks indicate significant difference between CK-MG and CK, DS-MG and DS, respectively (t test, n=5, *p<0.05, **p<0.01, ***p<0.001). One-way ANOVA was conducted using Duncan’s new multiple range test, and different letters above the bars indicate significant differences (p < 0.05).

### Global analysis of transcriptome on ear leaf with drought and MG effects

Taking into consideration of MG’s effect on delaying leaf senescence, SPAD value of ear leaf under DS was significantly higher than control from 12 DAI (Fig. S.1). The ear leaf in each treatment was selected and used for transcriptomic analysis at 12 DAI. This analysis aims to investigate the underlying mechanisms involved in postponing the onset of leaf senescence through exogenous MG under DS. Three differential expression genes (DEGs) pair-comparison were carried out, including DS vs CK, CK- MG vs CK, and DS-MG vs DS. A total of 920, 1664, and 1591 DEGs were filtered for each pair-comparison. Kyoto Encyclopedia of Genes and Genomes (KEGG) enrichment indicated that the DEGs of drought effect (DS vs CK) was mainly enriched in biosynthesis of amino acid, starch and sucrose metabolism, plant hormone signal transduction, plant pathogen interaction and carbon metabolism (Fig. 4, D). The DEGs of MG effects under well-watered (CK-MG vs CK) were mainly enriched in starch and sucrose metabolism, carbon metabolism, phenylpropanoid biosynthesis, plant hormone signal transduction, and plant pathogen interaction (Fig. 4B, D). Moreover, the DEGs of MG effects under DS (DS-MG vs DS) were mainly enriched in starch and sucrose metabolism, biosynthesis of amino acid, phenylpropanoid biosynthesis, plant hormone signal transduction, and plant pathogen interaction (Fig. 4C, D). Interestingly, the KEGG pathways enriched in drought effects were predominantly downregulated, while those enriched in MG effects under DS were predominantly upregulated. Specifically, further investigation will be conducted to explore the underlying mechanisms of delaying the onset of leaf senescence, focusing on the KEGG pathways associated with the effects of MG.

**Fig. 4.**
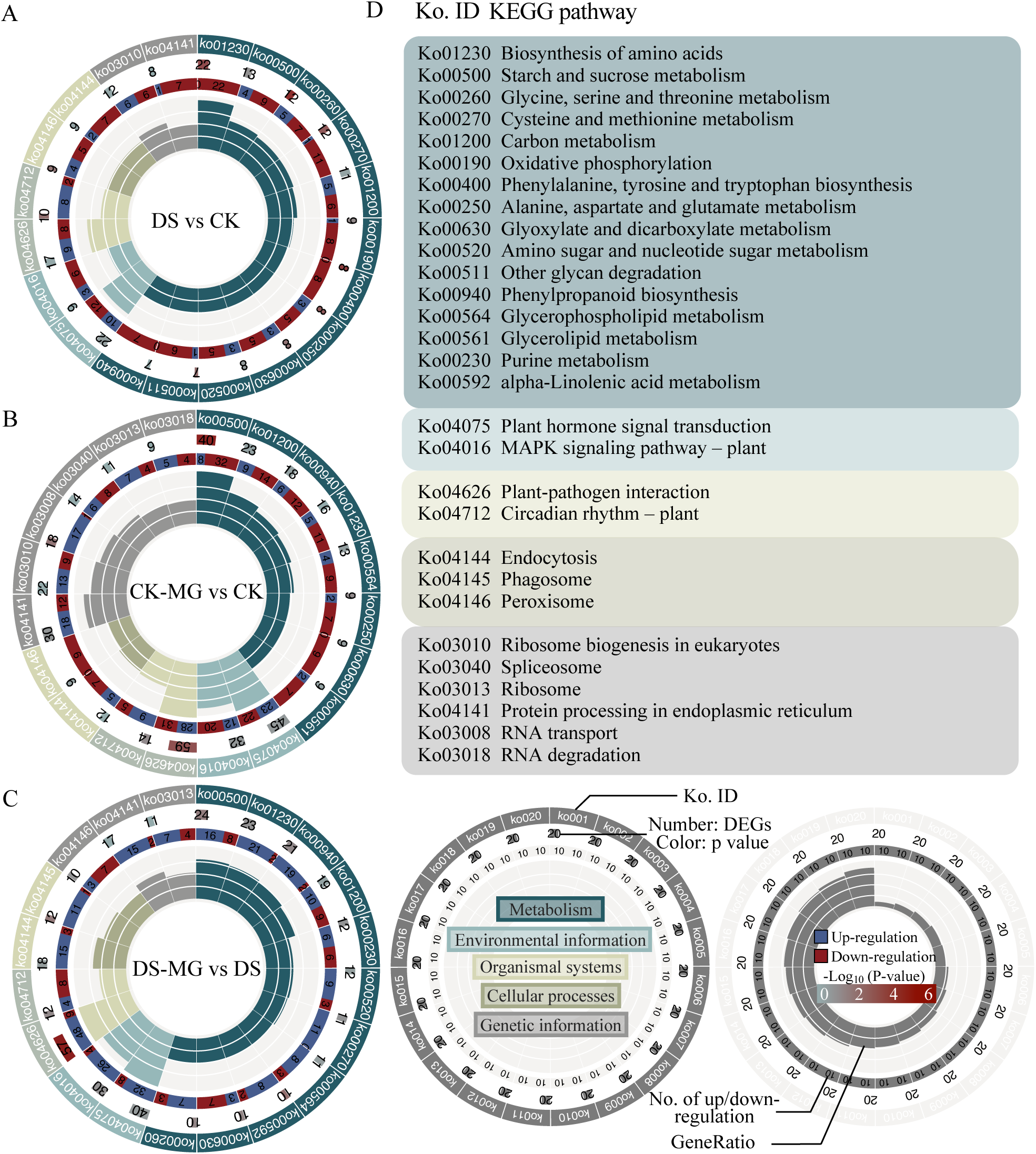
Global analysis of DEGs expression and KEGG pathway between (A) DS vs CK (drought effects); (B) CK-MG vs CK (MG effects under well-watered); and (C) DS- MG vs DS (MG effects under drought), respectively. For (A-C), the DEGs were classified into five categories, including metabolism (turquoise), environmental information (light turquoise), organismal systems (light khaki), cellular processes (khaki), and genetic information (gray). The KEGG pathway enrichment analysis of these top 20 DEGs was based on GeneRatio, and the corresponding KEGG pathways were listed in (D). The second circle represented the number of DEGs, and the third circle represented the number of up- and down-regulated DEGs in each KEGG pathway. The inner bar chart represented the normalized GeneRatio of each KEGG pathway. CK, well-watered; DS, drought stress.

### Inhibition of phloem loading and excessive accumulation of sugars under drought and MG effects

The enriched KEGG pathway in three pair-comparison have all associated to starch and sucrose metabolism. In total, 13, 40, and 24 of DEGs in drought effect and both MG effects under well-watered and DS were identified, respectively (Fig. 4A). In phloem loading for long-distance sugar transport, ZmSWEET13a, b, c facilitate the export of sucrose from parenchyma cells into the apoplast for subsequent uptake by ZmSUT1 from the apoplast into the sieve elements (Slewinski et al., 2009; Bezrutczyk et al., 2018). Under DS, ZmSWEET13a, b, c were down regulated, but ZmSUT1 was upregulated (Fig. 5A). Under CK-MG, ZmSWEET13a, b, c and ZmSUT1 were all downregulated (Fig. 5A). Similarly, under DS-MG, ZmSWEET13a, b and ZmSUT1 were downregulated but ZmSWEET13c was up regulated (Fig. 5A). These results indicate that the phloem loading was inhibited under drought and MG effects. Moreover, glucose content was increased by 21-108%, and sucrose content was increased by 24-30%, from 3-15 DAI under DS (Fig. 5B). The fructose content was increased by 22- 69% from 9-15 DAI under DS (Fig. 5B). Instead, starch content was decreased by 9- 33% from 9-12 DAI under DS (Fig. 5B). Similarly, under CK-MG, glucose content was increased by 60-160% from 6-15 DAI (Fig. 5C). Under CK-MG, fructose and sucrose content were increased by 11-99%, and 29-41% from 3-15 DAI, respectively (Fig. 5C). In contrast, starch content was decreased by 7-20% from 6-15 DAI under CK-MG (Fig. 5C). Under DS-MG glucose content was increased by 90-288%, and increased sucrose content by 50-78% from 9-15 DAI (Fig. 5D). Exogenous MG under DS increased fructose content by 38-126%, and decreased starch content by 5-25% from 6-15 DAI (Fig. 5D). The results reveal that drought and both MG effects caused sugar accumulation in leaf tissue. In addition, among the condition of sugar accumulation, family genes encoded trehalose phosphate synthase (TPS) were upregulated under DS and downregulated under CK-MG and DS-MG, while family genes encoding trehalose phosphate (TPP) were downregulated under DS and upregulated under CK- MG and DS-MG. These results suggest that trehalose-6-phosphate (T6P) may accumulate under DS but reduce under CK-MG and DS-MG.

**Fig. 5.**
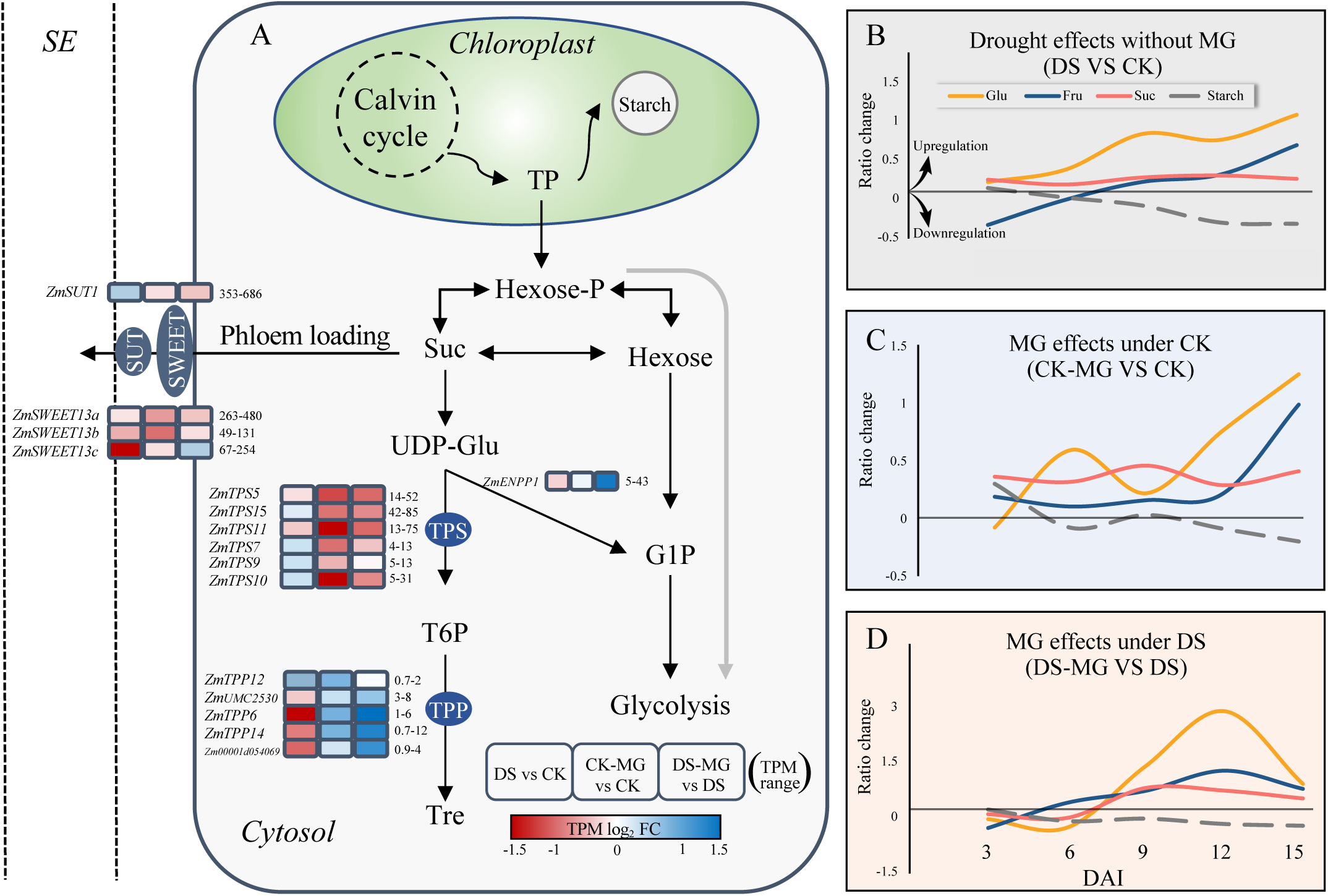
(A) Expression profile of sucrose transportation and T6P metabolism related gene and (B-D) the glucose, fructose, sucrose, and starch content in leaf under drought and MG effects, respectively. SE, sieve elements; CK, well-watered; DS, drought stress; DAI, days after drought induction.

### Exogenous MG induced glycolysis and glyoxalase system for maintaining endogenous MG levels

We further investigated the DEGs in glycolytic pathway under sugar accumulation by drought and both MG effects. The genes related to upstream of glycolysis, including phosphoglycerate mutase, fructose kinase, and phosphofructokinase (PFK) were upregulated under DS, CK-MG, and DS-MG (Fig. 6A). However, the genes related to downstream of glycolysis, including triosephosphate isomerase (TPI), glyceraldehyde-3-phosphate dehydrogenase (GAPDH), dihydrolipoamide dehydrogenase, pyruvate dehydrogenase E1 component alpha subunit, and pyruvate dehydrogenase E1 component beta subunit were downregulated under DS but upregulated under CK- MG and DS-MG (Fig. 6A). Additionally, PFK activity was significantly increased by 45% and GAPDH activity was decreased by 22% under DS (Fig. 6B, C). Conversely, exogenous MG significantly increased PFK and GAPDH activity by 42% and 26% and under well-watered, respectively (Fig. 6B, C). Similarly, exogenous MG significantly increased PFK and GAPDH activity by 11% and 82% under DS, respectively (Fig. 6B, C).

**Fig. 6.**
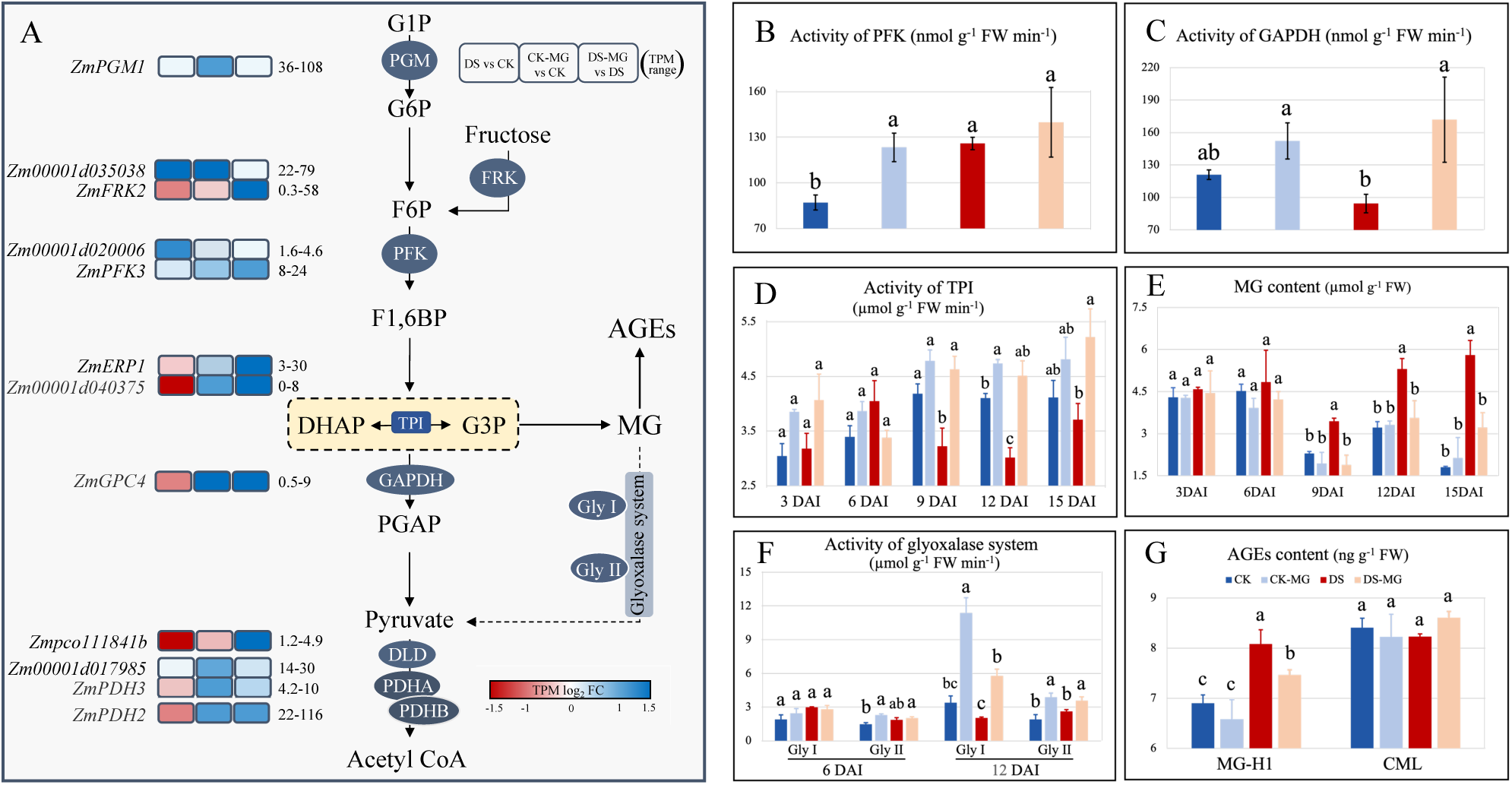
(A) Overview the expression pattern of glycolysis pathway, glycolytic enzyme activity and metabolite content under drought and MG effects, including details regarding (B, C) activity of PFK, GAPDH at 12 DAI, (D) TPI and (F) glyoxalase system, and the content of (E) MG and (G) AGEs at 12 DAI. CK, well-watered; DS, drought stress; DAI, days after drought induction; MG-H1, methylglyoxal hydroimidazolone; CML, carboxymethyl-lysine. Data are means (± SE). One-way ANOVA was conducted using Duncan’s new multiple range test, and different letters above the bars indicate significant differences (p < 0.05).

Endogenous MG was mainly originated from the precursor glyceraldehyde-3- phosphate and dihydroxyacetone phosphate which were regulated by the TPI. The activity of TPI under DS was significantly decreased by 10-26% compared to well- watered from 9-15 DAI (Fig. 6D). Instead, exogenous MG increased the activity of TPI by 14-27% under well-watered, by 28-50% under DS, respectively, from 3-15 DAI (except exogenous MG at 6 DAI under DS) (Fig. 6D). Correspondingly, MG level was significantly increased by 50-222% under DS from 9-15 DAI, while significantly decreased in MG level by 33-45% under DS-MG from 9-15 DAI (Fig. 6E). Furthermore, MG-scavenging system was determined to verify the potential detoxification capability under drought and both MG effects. At 6 DAI, DS, CK-MG, and DS-MG do not induce the activity of Gly I and Gly II (Fig. 6F). However, exogenous MG significantly increased Gly I activity by 230% under well-watered and increased by 182% under DS at 12 DAI, respectively (Fig. 6F). Exogenous MG significantly induced Gly II activity by 104% under well-watered and 36% under DS at 12 DAI, respectively (Fig. 6F). These results represent that MG effects reduced the potential for endogenous MG accumulation by increasing activity of TPI, Gly I, and Gly II. The AGEs MG-H1 adduct and CML adduct were investigated to represent the specific modification of protein by endogenous MG. MG-H1 adduct was significantly increased by 17% under DS, while MG-H1 adduct was significantly decreased by 8% under DS-MG. (Fig. 6G). Conversely, CML adduct was not have clear difference between each treatment (Fig. 6G).

In vitro, MG scavenger NAC reduced MG toxicity and maintained chlorophyll content under high sugar MG toxicity was increased in senescing leaf under DS but reduced under DS-MG (Fig. 6). In order to further verify whether MG toxicity was sufficient to cause leaf senescence under high sugar levels, an in vitro leaf strip senescence assay was performed (Fig. 7). Compared to water control, chlorophyll content was decreased by 10% under mannitol, 24-25% under sugars, and 40-55% under over-dosed MG application (Fig. 7). The declination of chlorophyll content was more prominent under hexoses or sucrose application than mannitol. This result indicates that leaf senescence was attributed to sugar effect excluded the factor of osmotic potential (Fig. 7). However, dose-depended manner of NAC (MG scavenger) application significantly maintained the chlorophyll content by 26% under glucose (1 mM), 13- 15% under fructose (1, 2 mM), 11-26% under sucrose, 26-81% under both over- dosed MG application, respectively (Fig. 7). These results indicate that MG toxicity resulting from excessive sugar content is qualified to cause leaf senescence and can be reversed by reducing MG toxicity with NAC application (Fig. 7).

**Fig. 7.**
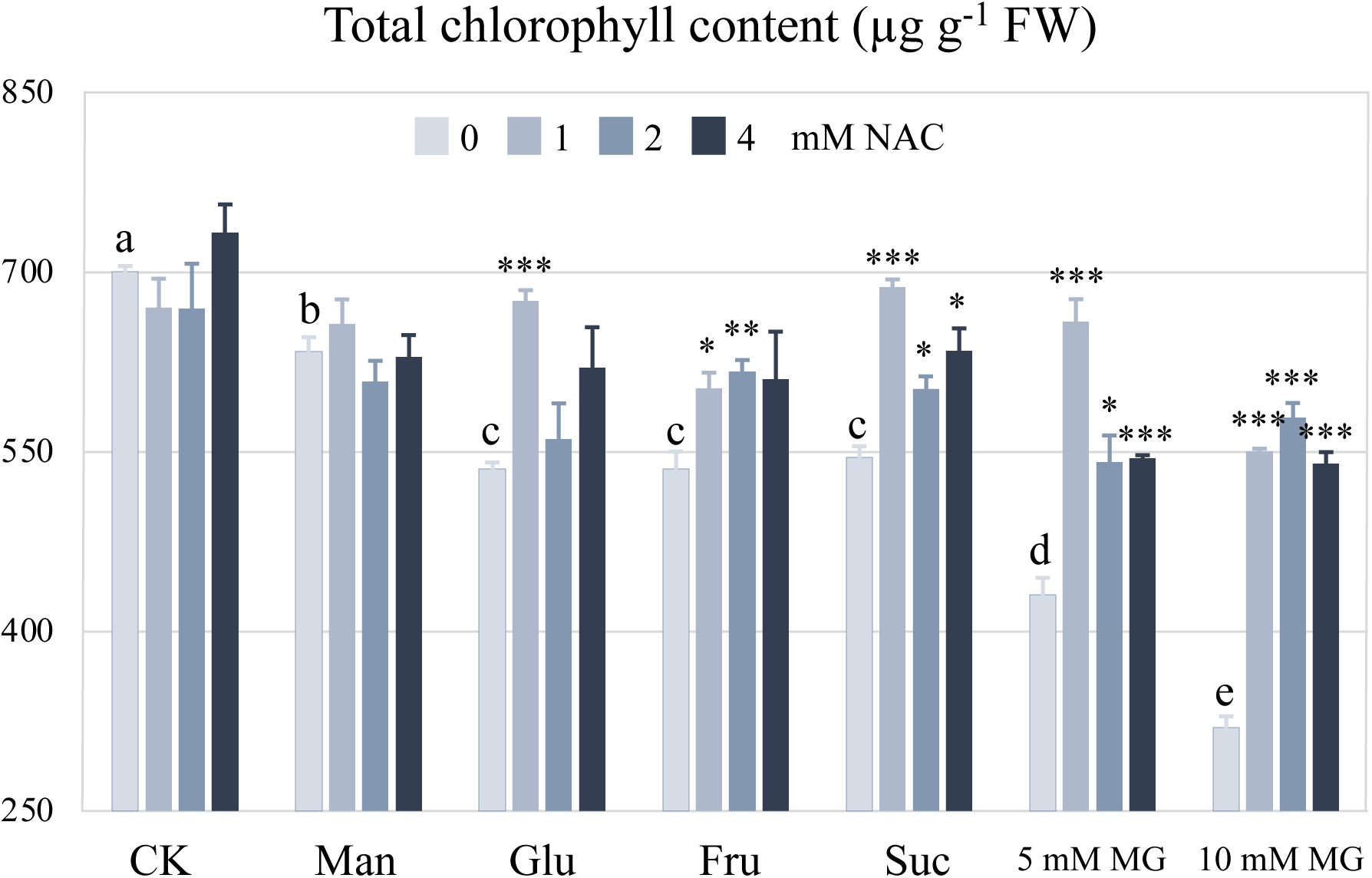
In vitro leaf strip senescence assay under distilled water (CK), mannitol (Man), glucose (Glu), fructose (Fru), sucrose (Suc), and MG solution with 0, 1, 2, 4 mM of N-acetyl-L-cysteine (NAC), respectively. Data are means (± SE). Asterisks indicate significant difference between NAC and without NAC application under each circumstance (t test, n=4, *p<0.05, **p<0.01, ***p<0.001). One-way ANOVA was conducted using Duncan’s new multiple range test, and different letters above the bars indicate significant differences (p < 0.05).

Exogenous MG stimulated phenylpropanoid biosynthetic pathway under drought The KEGG enriched analysis for DEGs were associated with phenylpropanoid pathway under CK-MG and DS-MG (Fig. 4B-D). Six enzymes involved in phenylpropanoid pathway were identified from DEGs and labeled in (Fig. 7A), including phenylalanine ammonia-lysase (PAL), phenylalanine/tyrosine ammonia-lysase (PTAL), caffeoyl-CoA O-methyltransferase (CCoAOMT), cinnamoyl-CoA reductase (CCR), cinnamyl-alcohol dehydrogenase (CAD), and peroxidase under drought and both MG effects (Fig. 8A). The genes related to the phenylpropanoid pathway were downregulated under DS but upregulated under DS-MG. Besides, CK-MG induced the genes related to the peroxidase in the phenylpropanoid pathway (Fig. 8B).

**Fig. 8.**
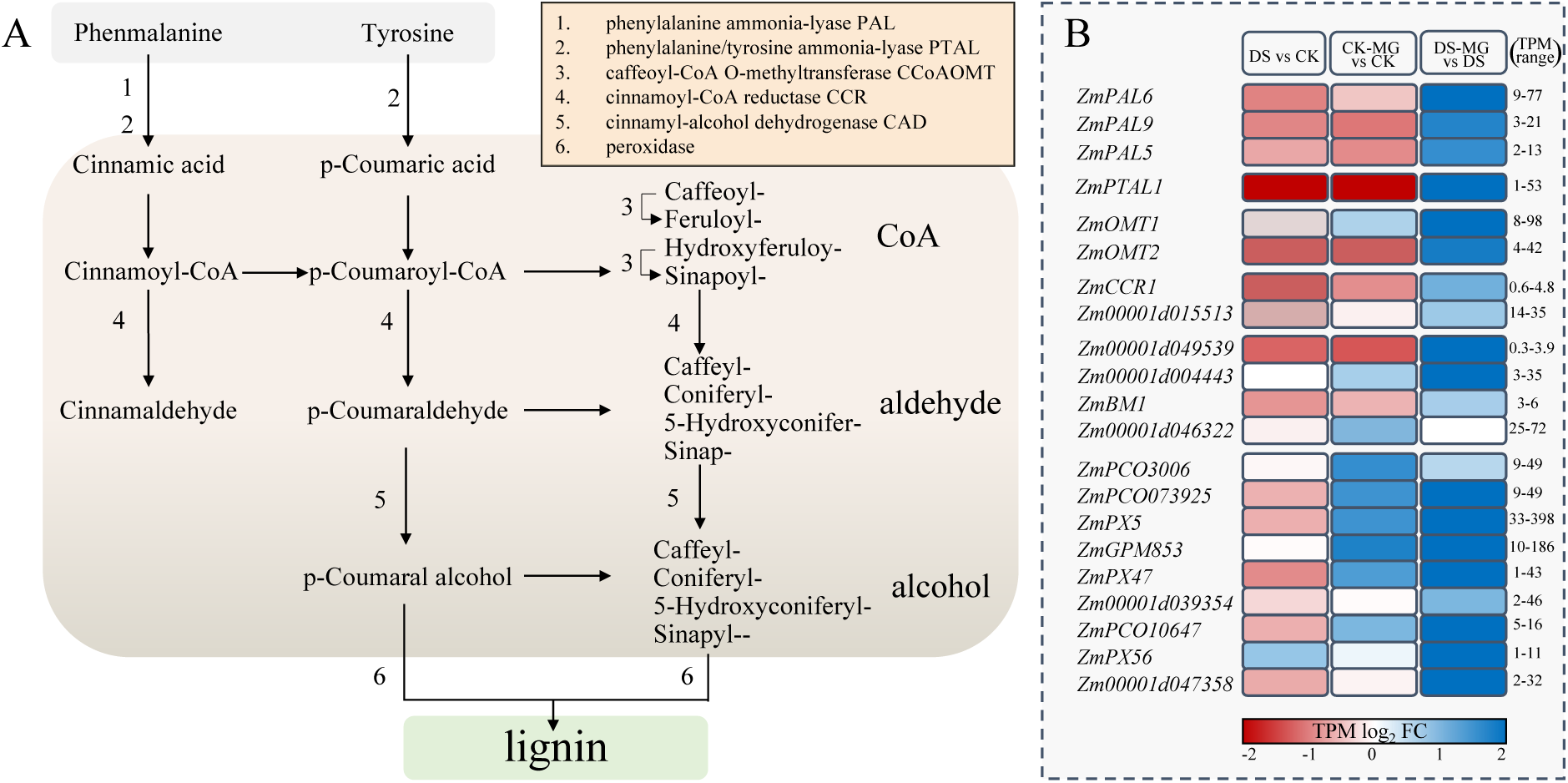
Overview of phenylpropanoid biosynthesis under drought and MG effects, including (A) catabolic location and (B) expression pattern of DEGs. CK, well- watered; DS, drought stress.

## Discussion

### Exogenous MG maintains source strength and retrieves yield penalty

Grain filling stage is particularly vulnerable to drought, as it inhibited source-sink communication and led to the severe yield losses (NeSmith and Ritchie, 1992; Lu et al., 2015; Barutcular et al., 2016). During this stage, kernel development relied on the considerable sugar from the ear part of leaves (Shen et al., 2020a; Shen et al., 2022; Shen et al., 2023). In this study, exogenous MG both increased kernel number and kernel weight, thus retrieving yield loss by 14% under DS; while had no significant effect on ear phenotype or yield performance under well-watered condition (Fig. 2; Fig. 9). Exogenous MG also delayed leaf senescence across the plant under DS and enhanced the photosynthetic capability under well-watered and DS (Fig. 3; Fig. 9). Similarly, previous studies demonstrated that enhancing the MG detoxification by overexpression of OsGly I gene in rice or double transgenic plants overexpressing Gly I and Gly II in tobacco increased yield output and stay-green phenotype under abiotic stresses (Singla-Pareek et al., 2003; Zeng et al., 2016). This evidence supports that increasing capability of MG detoxification is related to higher yield potential under adverse conditions. In this study, exogenous MG also induced the glyoxalase system, which efficiently reduced cytotoxic MG and developed the kernel normally under DS (Fig. 2; Fig. 6E, F). Interestingly, the sugar transporters facilitating phloem loading were suppressed by both DS and MG application (Fig. 5A), however, the yield output was significantly reduced under DS but maintained under DS-MG (Fig. 2E). We hypothesize that it could be because that (i) inhibited phloem loading and excessive sugar accumulation may not be sufficient to reduce yield, but the susceptibility to excessive sugar accumulation in the source leaf with or without MG application. (ii) Second, according to substrate concentration of sugar was higher than control under CK-MG and DS-MG. It still can transport enough photoassimilate to kernel even if the phloem loading was inhibited. (iii) Third, exogenous MG prolonged and improved the function of ear leaf during the drought may compensate for the inhibition of phloem loading. However, the underlying mechanisms in kernel deserve further investigation.

**Fig. 9.**
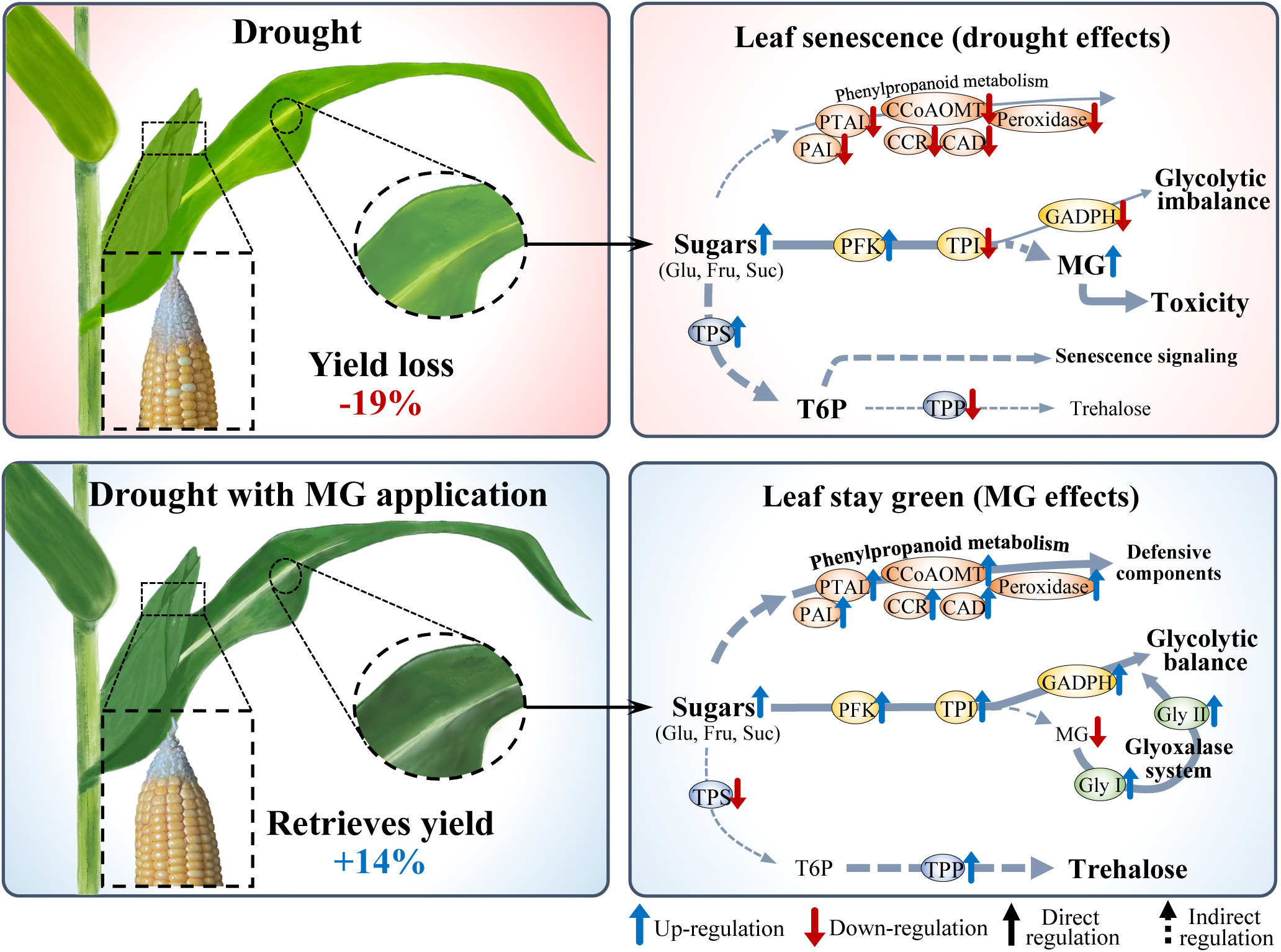
The model showing the effects of exogenous MG on leaf senescence, yield performance, and the underlying mechanisms involved in maintaining source strength under drought conditions. Glu, glucose; Fru, fructose; Suc, sucrose; PFK, phosphofructosekinase; TPI, triosephosphate isomerase; GAPDH, glyceraldehyde-3-phosphate dehydrogenase; Gly I, glyoxalase I; Gly II, glyoxalase II; PAL, phenylalanine ammonia-lysase; PTAL, phenylalanine/tyrosine ammonia- lysase; CCoAOMT, caffeoyl-CoA O-methyltransferase; CCR cinnamoyl-CoA reductase; CAD cinnamyl-alcohol dehydrogenase; TPS, phosphate synthase; TPP, trehalose phosphate; T6P, trehalose-6-phosphate.

### Exogenous MG does not alleviate drought-induced sugar accumulation but increases the resistance to high sugar

Progression of leaf senescence induced by abiotic stresses has been intensively discussed (Rolland et al., 2006; Janse van Rensburg et al., 2019). Sugar accumulation has been identified as a vital factor in the initiation of leaf senescence. For instance, exogenous sucrose and hexose have been found to induce rapid leaf senescence in tobacco and Arabidopsis, respectively, through sugar signaling by T6P and hexose (Wingler et al., 2004; Asim et al., 2022a). On the other hand, Arabidopsis mutant gin2 which lacking catalytic activity of hexokinase delays leaf senescence under low or high light intensity. This evidence indicates that the sugar signaling, and glycolytic sensitivity were associated with leaf senescence. However, the mechanism by which sugar accumulates in cell and regulates the leaf senescence under drought is still vague. In our study, sugar accumulation was prominent with the declination of starch content under drought and both MG effects (Fig. 5). We suggest that sugar accumulation was credited to the inhibition of phloem loading under drought and both MG effect, also the enhancement of photosynthetic capacity under both MG effects (Fig. 3D, E). Moreover, the product of photosynthesis possibly leans to transport out of the chloroplast rather than synthase to starch, causing the higher level of soluble sugar but lower level of starch (Fig. 3D, E; Fig. 5). Furthermore, T6P may accumulate under DS and decrease under CK-MG and DS-MG due to the transcriptional regulation of TPS and TPP (Fig. 5A). Previously, it has been confirmed that T6P is required in the onset of leaf senescence in Arabidopsis (Wingler et al., 2012), serving as an indicator of sucrose availability and negatively regulated SnRK1 (Janse van Rensburg et al., 2019). The mutant AtotsB lacking the activity of TPS delays leaf senescence, and has similar phenotype to overexpression of SnRK1 gene, KIN10 (Wingler et al., 2012). These results support that T6P regulation in leaf may cause the leaf senescence under drought but delay under MG effects through T6P signaling with T6P-SnRK1 interaction (Wingler et al., 2012) (Fig. 9).

High sugar content is coupled with excessive MG and AGEs accumulation which negatively affected leaf development in Arabidopsis seedlings (Borysiuk et al., 2018). The mutant pdtpi in Arabidopsis resulted in reduction of TPI activity and high-level MG accumulation, leading to severe stunting and chlorosis in seedling leaf. The author proposed that chloroplast morphology is defective in pdtpi possibly attributed to MG toxicity (Chen and Thelen, 2010). Nonetheless, the effects of MG and AGEs on leaf senescence have not been reported yet. In this study, exogenous sugars or over- dosed MG accelerated the leaf senescence but delayed by reducing MG toxicity with NAC application. This result indicates that MG toxicity is involved and qualified to cause leaf senescence under high sugar conditions (Fig. 7). Moreover, under the high sugar condition, upstream of glycolysis was upregulated under DS, CK-MG, and DS- MG (Fig. 6A-C). This result implies that glycolysis was rapidly responded to high sugar content. Remarkably, downstream of glycolysis was inhibited and started where the MG was formed under DS but upregulated under both MG effects (Fig. 6A, D; Fig. 9). These results imply that exogenous MG did not directly reduce the sugar content in cell but regulate the glycolytic flux in response to high sugar under DS. Additionally, exogenous MG induced higher level of TPI activity with lower level of endogenous MG and AGEs adduct under well-watered and DS (Fig. 6D, E, G). Specifically, both MG effects on lower MG toxicity to leaf cells may be the mutual consequence by increasing TPI activity and glyoxalase system (Fig. 6D-G; Fig. 9). According to the timeline of leaf senescence, sugar was rapidly accumulated from 3 DAI (Fig. 5B-D), TPI activity declined, and MG content rose from 9 DAI (Fig. 6 D, E), SPAD and photosynthetic capacity reduced from 12 DAI (Fig. 3). Moreover, the qualification of MG toxicity in relation to leaf senescence by in vitro leaf strip assay (Fig. 7). We suggest that sugar accumulation with disequilibrium in glycolysis caused the MG toxicity in cell and further accelerated leaf senescence under DS (Fig. 7). In contrast, exogenous MG increased higher sugar content while declining MG content by increasing glycolytic flux and glyoxalase system. Consequently, exogenous MG alleviates plant diabetes and ultimately led to the delay of leaf senescence under DS.

Exogenous MG activates phenylpropanoid biosynthesis under drought Phenylpropanoids pathway contributes to lot of aspects of responses toward environmental stimuli and within vast variety of antioxidant phenolics or produced the defensive component (Fraser and Chapple, 2011; Wang et al., 2021). Specifically, the downstream of phenylpropanoid pathway involves lignin biosynthesis, which is essential for structural integrity and pathogen resistance (Vogt, 2010; Chan et al., 2023). Previously, it was confirmed that transcription factor MeRAV5 promoted the MeCAD15 to affect lignin accumulation which increased drought tolerance in cassava (Yan et al., 2021). In this study, exogenous MG upregulated the genes involved in phenylpropanoid pathway including PAL, PTAL, CCoAOMT, CCR, CAD, and peroxidase under DS (Fig. 8; Fig. 9). These results imply that exogenous MG may positively regulate the phenolic metabolites and lignin accumulation to increase drought tolerance. Interestingly, the lodging rate was reduced under CK-MG and DS-MG at 65 DAP (Fig. S.2). It supports the effect of exogenous MG on lignin biosynthesis across the plant and maintaining the higher structural integrity under well-watered and DS. It has been characterized that phenylpropanoid metabolism is an irreversibly carbon-energy consuming process that requires around 30-40% of all organic carbon sources (Van Heerden et al., 1996). Additionally, lignin possesses a relatively higher energy density than cellulose and hemicellulose by 30% (Novaes et al., 2010). Therefore, we speculate that exogenous MG significantly upregulated the phenylpropanoid metabolism possibly due to the sufficient carbon source with sound energy production by the balance of glycolysis under DS (Fig. 6A; Fig. 7).

## Conclusion

Drought causes an accumulation of sugars in leaves by inhibiting phloem loading. Consequently, this disruption in sugar metabolism causes an imbalance in glycolysis and results in excessive accumulation of endogenous MG, which induces leaf senescence. Conversely, exogenous MG confers drought tolerance with alleviation of leaf senescence and yield loss by stimulation of glycolytic flux and glyoxalase system to enhance the bearing capability to high sugar. Besides, exogenous MG induces the phenylpropanoid biosynthetic pathway, which may associate with modulation of drought tolerance. Collectively, these findings hold both theoretical and practical significance to maintain yield in maize production in face of drought stress.

## Acknowledgments

This work was supported by the the National Key Research and Development Program of China (2022YFD2300803; 2022YFD1900702), National Natural Science Foundation of China (32272013), the China Agriculture Research System (CARS-02-16).

## Author Contribution

Y.-H. L., S. S., and S.-L. Z. designed the study; Y.-H. L., Y.-N. Z., Y.-K. J., Z.-D. X., Y.-J. Z, C. H., B. H., and Z.-Y. C. performed the experiments and collected the data; Y.-H. L., Y.-J. Z., C. H. analyzed and interpreted the data; Y.-H. L., S. S., and S.-L. Z. wrote the drafts; Y.-H. L., S. S., and S.-L. Z. revised the manuscript. All authors approved the final version of the manuscript.

## Data availability

All relevant data can be found within the manuscript and its supporting meterials. The sequencing data will be available at NCBI.

